# Respiratory viral infection alters the gut microbiota by inducing inappetence

**DOI:** 10.1101/666354

**Authors:** Helen T. Groves, Sophie L. Higham, Miriam F. Moffatt, Michael J. Cox, John S. Tregoning

## Abstract

The gut microbiota has an important role in health and disease. Respiratory viral infections are extremely common but their impact on the composition and function of the gut microbiota is poorly understood. We previously observed a significant change in the gut microbiota after viral lung infection. Here we show that weight loss during Respiratory Syncytial Virus (RSV) or influenza virus infection was due to decreased food consumption, and that fasting mice independently of infection altered gut microbiota composition. While the acute phase TNF-α response drove early weight loss and inappetence during RSV infection, this was not sufficient to induce changes in the gut microbiota. However, depleting CD8^+^ cells increased food intake and prevented weight loss resulting in a reversal of the gut microbiota changes normally observed during RSV infection. Viral infection also led to changes in the faecal gut metabolome during RSV infection, with a significant shift in lipid metabolism. Sphingolipids, poly-unsaturated fatty acids (PUFAs) and the short-chain fatty acid (SCFA) valerate all increased in abundance in the faecal metabolome following RSV infection. Whether this, and the impact of infection-induced anorexia on the gut microbiota, are part of a protective, anti-inflammatory response during respiratory viral infections remains to be determined.

## Introduction

The gut microbiota plays many critical roles in maintaining human health. These include local effects such as metabolising non-digestible nutrients, providing colonisation resistance against gut infection, helping maintain intestinal barrier function and educating the immune system (Sekirov *et al*., 2010). It is increasingly appreciated that the gut microbiota also has systemic effects on health, for example through the production of anti-inflammatory metabolites such as short-chain fatty acids (SCFAs)(Tan *et al*., 2014). In the context of respiratory disease, most studies have focused on how the gut microbiota influences immune responses in the airways (Hauptmann and Schaible, 2016). Changes in gut microbiota composition can change the gut metabolome, with a subsequent impact on host immune function (Corrêa-Oliveira *et al*., 2016). Feeding mice a diet high in fermentable fibre decreases lung damage and increases survival during influenza virus infection by increasing faecal and serum SCFA (Trompette *et al*., 2018). Likewise increased abundance of the poly unsaturated fatty acid (PUFA) docosahexanoic acid (DHA) after *Lactobacillus* supplementation led to reduced lung inflammation and damage during RSV infection in mice (Fujimura *et al*., 2014).

However, several studies have demonstrated that respiratory infections are associated with a change in the composition of the gut microbiota (Wang *et al*., 2014; Deriu *et al*., 2016; Bartley *et al*., 2017; Groves *et al*., 2018; Hanada *et al*., 2018; Yildiz *et al*., 2018). We have previously observed that viral lung infections alter the gut microbiota, leading to an increase in the relative abundance of *Bacteroidetes* and a decrease in the relative abundance of *Firmicutes* (Groves *et al*., 2018). In our previous studies, we did not identify a mechanism that linked viral lung infection with changes in the gut microbiota. We did note that changes in overall gut microbiota composition after either Respiratory Syncytial Virus (RSV) or influenza A virus infection were similar, implying that the underlying mechanism is common to both infections and therefore not a pathogen specific immune effect as suggested elsewhere (Wang *et al*., 2014; Deriu *et al*., 2016). In mice one common symptom after RSV or influenza infection is weight loss, part of a wider pattern of sickness behaviours (Langhans, 2000). This weight loss has been associated with reduced food intake after influenza infection in mice (Swiergiel, Smagin and Dunn, 1997; Wang *et al*., 2016). Though the effect of RSV infection on food intake in mice has not been published, mild anorexia was observed in RSV infected pre-term lambs (Meyerholz *et al*., 2004). Loss of appetite is also reported after human influenza virus and RSV infection (Monto *et al*., 2000; Widmer *et al*., 2012). Reduced calorie intake in humans and mice has been associated with a significant increase in the ratio of *Bacteroidetes* to *Firmicutes* abundance (Ley *et al*., 2006; Fabbiano *et al*., 2018; Wang *et al*., 2018), similar to that observed in our prior viral infection study. This suggests that changes in the gut microbiota seen after respiratory viral infection might be driven by reduced food intake.

Therefore the aim of this study was to understand the interplay between infection, food intake, the gut microbiome and the gut metabolome. We show that changes in the gut microbiome after infection are caused by reduced food intake and that this is associated with CD8^+^ T cells. We also observe a significant change in the gut metabolome after lung infection with a significant increase in the levels of lipids produced.

## Results

### Respiratory infection reduces food consumption and alters the gut microbiota

We first investigated the link between weight loss and food intake after infection. Mice infected with RSV lost weight from the first day after infection, stabilising between days 1 and 4 and then losing more weight resulting in approximately 15-20% weight loss by day 7 (Fig.1A). There was no weight loss in PBS dosed or naïve mice. Food consumption mirrored weight loss. The average amount of food consumed by one uninfected mouse was 3 g (±0.3) a day. RSV infected mice immediately ate less following infection (D0-D1: 1.3 g). Food consumption in RSV infected mice increased from day 1 to 4. After day 4, RSV infected mice ate less food each day, with the nadir food consumption of 0.5 g per mouse on day 6 (Fig.1B). We saw a similar effect after influenza infection (Fig.S1). This strongly suggests that respiratory viral infection induces inappetence which then reduces food consumption leading to weight loss.

**Figure 1:**
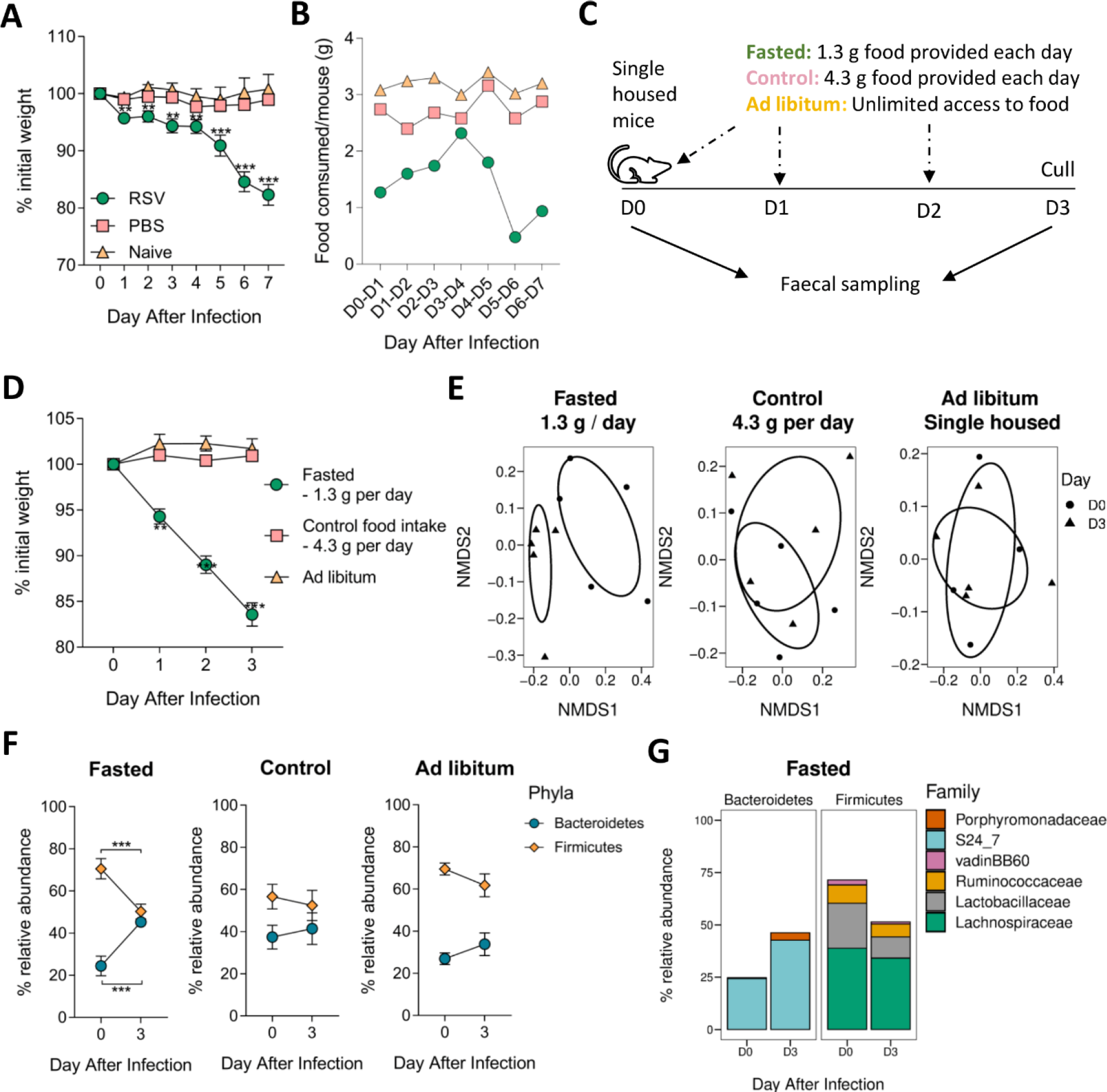
Reduced food consumption alters the composition of the gut microbiota and increases the relative abundance of Bacteroidetes and decreases Firmicutes. Mice were intranasally infected with RSV, or intranasally dosed with sterile PBS or left naïve. Weight of individual mice **(A)** and food consumption of the entire cage was measured every day **(B)**. Fasted mice were provided with 1.3 g food per day per mouse which was 30% of control food intake (4.3 g per mouse per day). Faecal samples were taken from fasted and control mice and from mice with unlimited access to food to control (*ad libitum)* at day 0 and day 3 **(C)**. Weight during fasting **(D)**. Fasted mice had a significantly different gut microbiota diversity composition compared to pre-fasting (*P* = 0.01) **(E)**. Fasted mice had an increase in the relative abundance of Bacteroidetes and decreased relative abundance of Firmicutes **(F)** and significantly higher relative abundance of the S24_7 family (*P* = 0.001) in their gut microbiota compared to control mice **(F)**. There was no difference in the gut microbiota composition of control mice or ad libitum mice during the experiment **(E-F)**. N = 5 mice per group. * *P* ≤ 0.05, ** *P* ≤ 0.01 *** *P* ≤ 0.001

Having observed that reduced food intake during infection was associated with weight loss, we wanted to investigate whether there was a link between food intake, weight loss and changes in gut microbiota outside the context of infection. To do this, we restricted access to food to reflect the weight loss seen after infection (Fig.1C). Single housing of mice was required to control food intake per mouse. But as mice are social animals, the stress of individual housing might impact the gut microbiota. To control for this, a cage of single-housed mice with *ad libitum* food access was monitored alongside *ad libitum* co-housed animals to account for any single housing, stress-induced changes. No differences in food consumption or gut microbiota diversity were seen between *ad libitum* individually housed or co-housed mice (Fig.S2). Fasted mice (with access to 1.3 g food per day) lost approximately 15% total weight by day 3 (Fig.1D). Neither the control mice receiving 4.3 g food per day nor the *ad libitum* single housed mice lost weight.

Faeces were collected at day 0 and 3 to sample the gut microbiota. Beta diversity was significantly different following fasting, indicating a shift in overall gut microbiota composition (*P*<0.01, Fig.1E). There was no difference in beta diversity between days 0 and 3 in control mice or single housed *ad libitum* mice. At the phylum level, the gut microbiota of fasted mice had a significantly higher relative abundance of Bacteroidetes and a significantly lower relative abundance of Firmicutes compared to baseline (Fig.1F). There was no change in these phyla in control or single housed *ad libitum* fed mice. At the family level, the major change seen was an increase in the abundance of the S24_7 family (also known as Muribaculaceae (Lagkouvardos *et al*., 2019): Fig.1G). This pattern was very similar to that previously observed after RSV infection(Groves *et al*., 2018).

### TNF-α is associated with weight loss after infection, but not changes in the gut microbiota

Having observed that viral infection reduces food intake and reducing food intake alters the gut microbiota, we wanted to determine the role of host immune factors in inappetence after respiratory viral infection. Injection of recombinant murine TNF-α (r-TNF-α) has been shown to induce weight loss in mice (Biesmans *et al*., 2015). TNF-α is elevated in the airways in response to RSV infection in both humans and mice (Jafri *et al*., 2004; McNamara *et al*., 2004) and blocking TNF-α during RSV infection has been shown to reduce weight loss (Rutigliano and Graham, 2004; Tregoning *et al*., 2010). Therefore, we examined the role of TNF-α on inappetence, food intake and the gut microbiota after RSV infection.

Mice were injected intraperitoneally with an anti-TNF-α monoclonal or a control antibody on days 0, 2, 4 and 6 of infection (Tregoning *et al*., 2010) with faecal samples taken on days 0, 3 and 7 of infection. Anti-TNF-α treated mice did not lose any weight during the first 5 days of RSV infection, in contrast to the isotype control group which lost weight immediately following infection. However, after five days, anti-TNF-α treated mice lost weight rapidly and by day 7 there was no difference in weight loss between them and the control group (Fig.2A). Infected isotype control mice reduced their average food intake between day 1 and 2 of infection (Fig.2B), whereas blocking TNF-α prevented inappetence during the early stages of infection (days 1 – 4), but from day 5 onwards TNF-α blocked mice had reduced food intake. Anti-TNF-α treated mice had a significantly higher RSV lung viral load at day 7 compared to the isotype control group (Fig.2C). There was no effect of TNF blockade on the T cell response (Fig.S3). Beta diversity corresponded with weight loss during RSV infection. RSV infected isotype control mice experienced a significant shift in gut microbiota beta diversity at both days 3 and 7, while TNF-α depleted mice only had a significantly altered gut microbiota beta diversity at day 7 (Fig.2D). At day 7 the change in the relative abundance of Bacteroidetes and Firmicutes during RSV infection was reduced in anti-TNF-α treated compared to the control group (Fig.2E). Anti-TNF-α treated mice had significantly less Ruminococcaceae and Lactobacillaceae in their gut microbiota at day 7 than baseline, whereas isotype control mice had fewer Lachnospiraceae and more S24_7 (Fig.2F).

**Figure 2.**
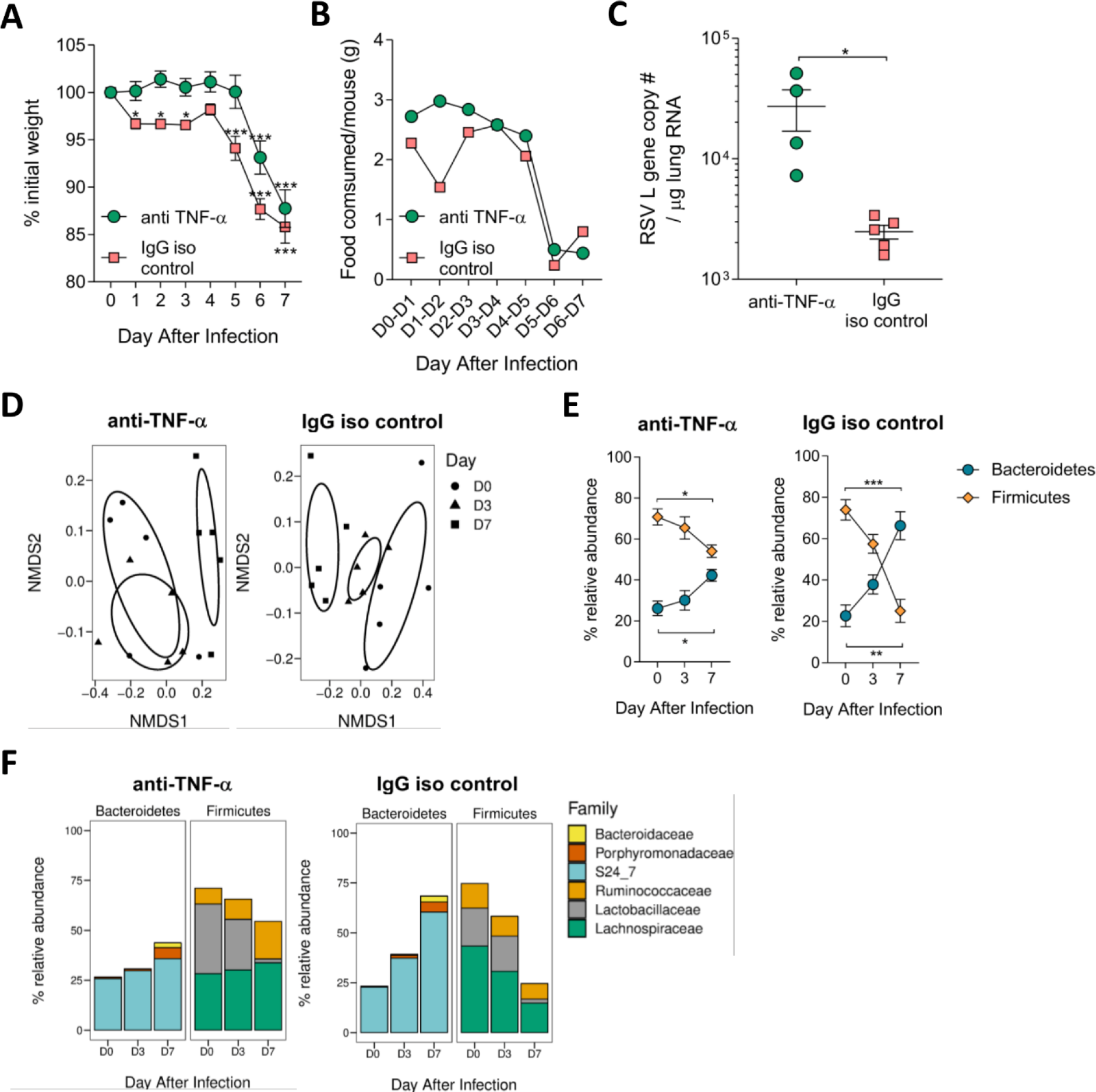
TNF-α drives early weight loss during RSV. Mice were IP injected with 450 µg/ml anti-TNF-α antibody or IgG isotype control before (-4hr) and after (D2, D4, D6) RSV infection. Faecal samples were taken before infection (D0) and at days 3 and 7. Weight loss was measured every day **(A)** as was food consumption **(B)**. RSV viral load in the lungs (day 7) was significantly higher in TNF-α depleted mice **(C)**. There was no difference in gut microbiota diversity at day 3 in TNF-α depleted mice, with a significant shift in beta diversity at day 7 (*P* = 0.01). Mice treated with isotype control had a significant difference in microbiota diversity at day 3 (*P* = 0.05) and 7 (*P* = 0.07) **(D)**. Depleting TNF-α still resulted in a significant increase in the relative abundance of Bacteroidetes and decrease in Firmicutes at day 7 post infection compared to day 0 **(E)**. At the family level there was no significant change in relative abundance at day 3 in either TNF-α depleted mice or the isotype control group, and a significant (*P* ≤ 0.05) decrease in the relative abundance of Ruminococcaceae and Lactobacillaceae at day 7 in TNF-α depleted mice and a decrease in Lachnospiraceae (*P* ≤ 0.05) and increase in S24_7 (*P* ≤ 0.0001) in the isotype control group **(F)**. N = 5 mice per group. Results representative of two experiments.

Since we saw a smaller change in gut microbiota in anti-TNF-α treated mice following RSV infection, we investigated whether airway TNF-α could alter the gut microbiota in the absence of infection. Based on the doses and subsequent weight loss reported after intraperitoneal TNF administration by Biesmans *et al*., (Biesmans *et al*., 2015), we intranasally delivered 3 µg recombinant-TNF-α (r-TNF-α) per mouse, aiming for 1 g weight loss every 24 hours. Mice intranasally dosed with r-TNF-α began losing weight on day 1 with peak weight loss on day 2. Mice began to recover slightly by day 3, although there was still significant weight loss compared to baseline (*P*<0.05, Fig.3A). Food intake mirrored weight loss: r-TNF-α dosed mice ate significantly less between 1 and 2 days after infection (Fig.3B). They began to eat more on day 3 despite receiving another dose of r-TNF-α. TNF-α levels in the airways of r-TNF-α dosed mice were significantly higher than in PBS mice (*P*<0.05, Fig.3C).

**Figure 3.**
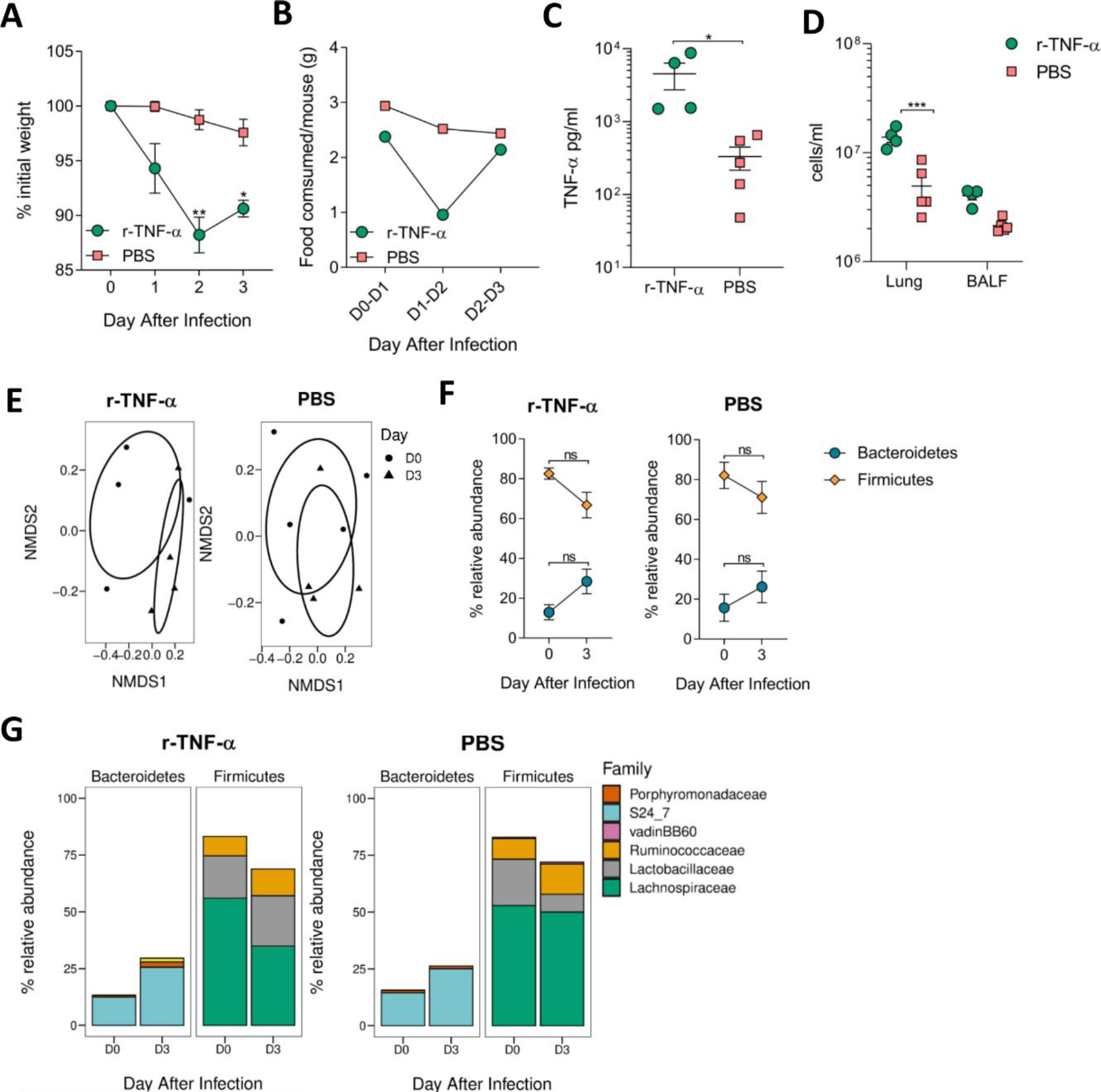
Acutely increasing airway TNF-α drives reduced food consumption and weight loss but is not sufficient to alter the gut microbiota composition. Levels of airway TNF-α were increased by intranasally dosing mice with 3 µg recombinant TNF-α or PBS every day for three days. Faecal samples were taken before (day 0) and after (day 3). This resulted in significant weight loss and reduced food intake **(A-B)**. Intranasally dosing mice with r-TNF-α did significantly increase airway TNF-α levels **(C)** and increased the total number of cells in the lungs **(D)**. There was no difference in the diversity of the gut microbiota after r-TNF-α or PBS dosing **(E)**. There was also no difference in the relative abundances of Bacteroidetes or Firmicutes **(F)** or in the abundance of individual bacterial families **(G)**. N = 4-5 mice per group. Results representative of two experiments.

r-TNF-α dosed mice had significantly higher total lung cell counts compared to PBS dosed mice (*P*<0.001, Fig.3D). However, while it induced acute weight loss, increasing the levels of airway TNF-α had no effect on gut microbiota beta diversity (Fig.3E). Both r-TNF-α and PBS dosed mice showed a trend towards increased relative abundance of Bacteroidetes and decreased relative abundance of Firmicutes (Fig.3F). There was no significant change in the abundance of families belonging to either phyla (Fig.3G). This data indicates that TNF-α is not the sole driver behind inappetence and weight loss during RSV infection.

### Depleting CD8^+^ T cells during RSV infection reduces inappetence and reverses changes in the gut microbiota

Blocking CD8^+^ T cells during RSV infection has been shown previously to reduce weight loss and increase viral load (Graham *et al*., 1991; Tregoning *et al*., 2008), and adoptive transfer of RSV-specific CD8^+^ T cells into naïve mice prior to RSV infection increases weight loss (Alwan, Kozlowska and Openshaw, 1994). We therefore investigated whether CD8^+^ T cells have a role in inappetence and the changes observed in the gut microbiota after infection.

CD8^+^ cells were depleted by systemic injection of an anti-CD8^+^ monoclonal antibody on days -1, 2 and 5 of infection. Control mice were injected at the same time with an IgG isotype control antibody. There was no effect on early weight loss, however depleting CD8^+^ cells prevented all subsequent weight loss compared to control mice (*P*<0.01, Fig.4A). Depleting CD8^+^ cells reduced, but did not completely prevent, inappetence (Fig.4B): anti-CD8 treated mice only ate 1.8 – 2 g of food per day over the last two days of infection compared to the usual 3 g/mouse/day. This was higher than the infected control group which were consuming an average of 0.3-0.5 g food per mouse on day 6 and 7. Viral load in the lungs significantly increased following CD8^+^ cell depletion (Fig.4C). CD8^+^ depletion significantly reduced the numbers of CD8^+^ T cells in the lungs after infection but not the overall number of cells recruited to the airways (Fig.S4).

**Figure 4.**
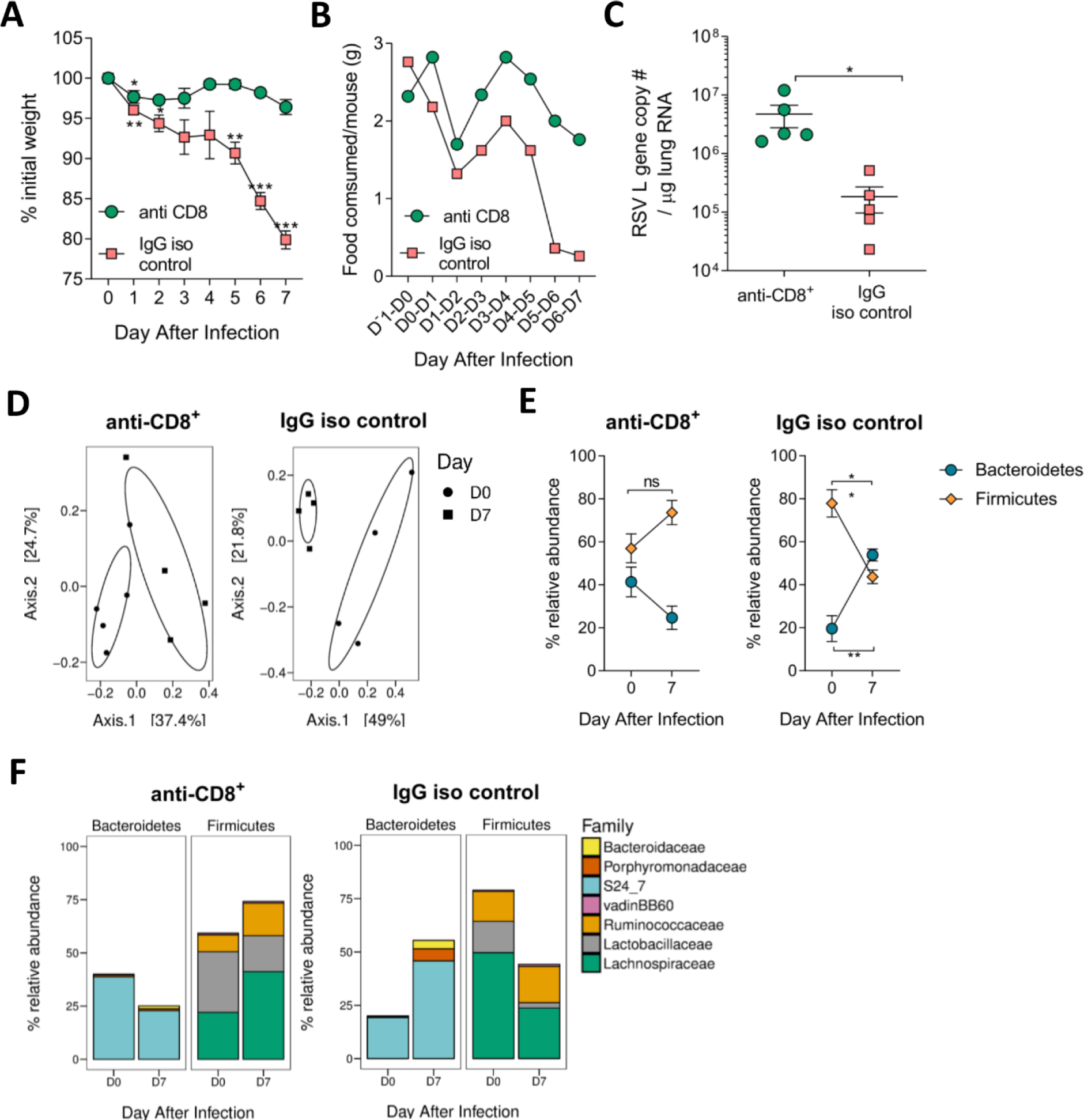
Blocking CD8+ T cells during RSV infection prevents weight loss and associated gut microbiota changes. CD8 T cells were blocked during RSV infection using mouse anti-CD8^+^ monoclonal antibody. Mice were IP injected with using 500 µg/ml antibody or isotype control before (Day -1) and after (D2, D5) RSV infection. Faecal samples were taken at D0 and D7. Blocking CD8^+^ T cells prevented weight loss at D2-D7 post infection compared to D0 **(A)** and reduced, but did not completely prevent, infection-induced anorexia **(B)**. The overall gut microbiota composition was still significantly different post RSV infection, in both anti-CD8^+^ treated mice (*P* = 0.04) and in the control group (*P* = 0.03) **(C)**. There was an increase in the relative abundance of Bacteroidetes and a decrease in Firmicutes in the control mice but no change in the ratio of these phyla in the anti-CD8^+^ treated mice **(D)**. At the family level, there was a significant decrease in the relative abundance of the S24_7 family (*P* ≤ 0.05) and increase in Lachnospiraceae (*P* ≤ 0.01) when CD8^+^ T cells were blocked during RSV infection, whereas the opposite was seen in the isotype control group (S24_7 increase, *P* ≤ 0.01 and Lachnospiraceae decrease, *P* ≤ 0.01) **(E)**. N = 4-5 mice per group.

Despite anti-CD8 treatment preventing weight loss during infection, there was a significant shift in gut microbiota beta diversity from day 0 to 7 (Fig.4D). Interestingly there was no change in the relative abundance of Bacteroidetes or Firmicutes following CD8^+^ cell depletion, unlike the control group (Fig.4E). Blocking CD8^+^ cells reversed the effect of RSV infection on the bacterial families detected in the gut, with a significant decrease in the relative abundance of S24_7 and an increase in Lachnospiraceae (Fig.4F). Depleting CD8^+^ cells during RSV infection reduced inappetence and immediate weight loss, with a marked effect on the gut microbiota.

### RSV infection changes the faecal metabolome, altering lipid metabolism

We hypothesized that reduced food intake during infection would decrease overall nutritional availability within the gut, impacting host and microbiota metabolism and metabolite levels. The faecal metabolome has been shown to be altered by both acute and long-term fasting (Rubio-Aliaga *et al*., 2011; Zheng *et al*., 2016; Collet *et al*., 2017). This is likely to be a combination of direct effects of fasting on host metabolism and indirect effects of altered gut microbiota composition and metabolism.

Faecal metabolomics was used to assess how gut metabolism changed during RSV infection. Mice were intranasally infected with RSV and faecal samples were taken every day for 7 days. The samples were processed, analysed by mass spectrometry and annotated(Evans *et al*., 2014; Farne *et al*., 2018). There were 803 detected biochemicals, 705 of which could be identified. The overall faecal metabolomic profile significantly shifted over time following RSV infection (*P*=0.02, Fig.5A). When comparing each time point to baseline, days 3, 6 and 7 had a significantly different faecal metabolic composition, which coincided with when the most significant weight loss occurred (Fig.S5A).

**Figure 5:**
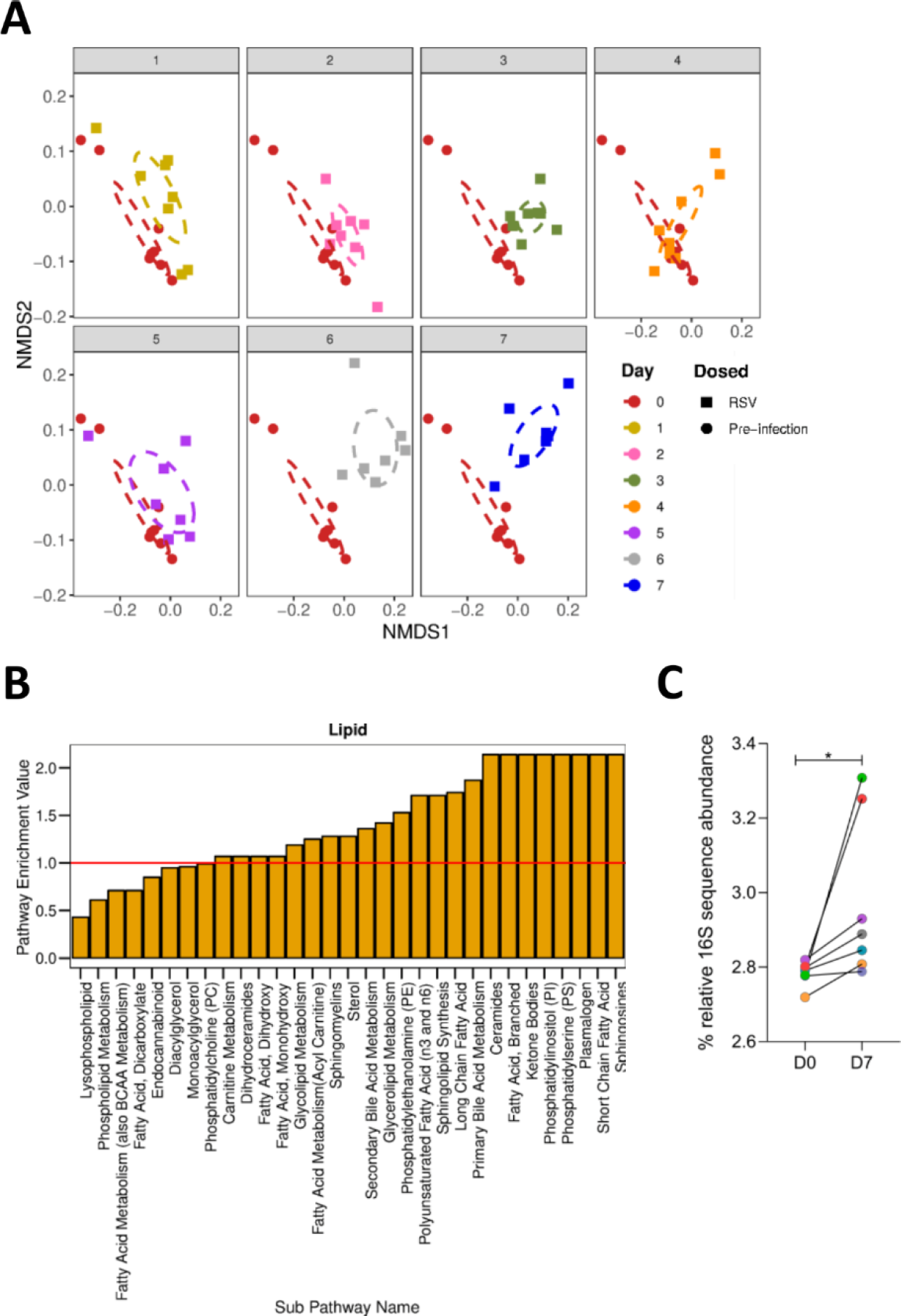
The gut metabolome, and particularly lipid metabolism, changes during RSV infection and is associated with predicted changes in the functional capacity of the gut microbiota. Mice were infected with RSV and faecal samples were taken before infection (D0) and every day post infection until day 7 (D7). N = 7-8 mice. The faecal metabolome was profiled using HPLC-MS. The overall composition of the faecal metabolome shifted significantly during RSV infection (*P* = 0.002) with different profiles detected at day 3 (*P* = 0.006), day 6 (*P* = 0.006) and most significantly at day 7 (*P* = 0.001) **(A)**. Pathway enrichment analysis examining sub-metabolic pathways (x axis) belonging to the lipid metabolism super pathway, which contain significantly altered abundances of metabolites after RSV infection (day 7), compared to before infection. A pathway enrichment value of >1 (red line) indicates that this pathway contains more experimentally different metabolites relative to the study as a whole. **(B)**. Predictive metagenomic and functional changes using PICRUSt on 16S rRNA sequencing data revealed a significant change in bacterial gene sets belonging to 7 KEGG orthology pathways, one of which was lipid metabolism (*P* = 0.02) **(C)**. PICRUSt analysis of corresponding 16S rRNA gene sequencing data with relative sequence abundance of predicted KEGG orthologs classified as belonging to lipid metabolism pathways (colours correspond to individual mice, lines link paired samples) **(C)**.

Seventy-six sub-pathways contained biochemicals which were significantly altered in abundance at day 7 compared to day 0 (Fig.S6). Fifty-three of these pathways had a pathway enrichment value greater than 1, and 23 of these belonged to the lipid metabolism super-pathway (Fig.5B). As seen previously, RSV infection significantly altered the gut microbiota (Fig.S5B-D). To link changes seen in the metabolome with the changes in the gut microbiota we used PICRUSt to perform predictive metagenomic analysis. There was a significant increase in the relative abundance of 16S rRNA gene sequences with predicted orthology functions in lipid metabolism at day 7 after RSV infection (*P*<0.05, Fig.5C). This reflected the observed change in the faecal metabolome suggesting that changes in lipid metabolism were driven by changed gut microbiota.

The sphingolipid and fatty acid metabolism pathways had some of the highest pathway enrichment values following RSV infection. Metabolites within the sphingosines, sphingolipids, sphingomyelins and ceramide sub-pathways were significantly increased in abundance at day 7 after RSV infection compared to baseline (Fig.6A). Multiple PUFAs were increased in abundance following RSV infection, including the anti-inflammatory DHA. The SCFA valerate was also significantly increased in abundance following RSV infection (Fig.6A).

**Figure 6:**
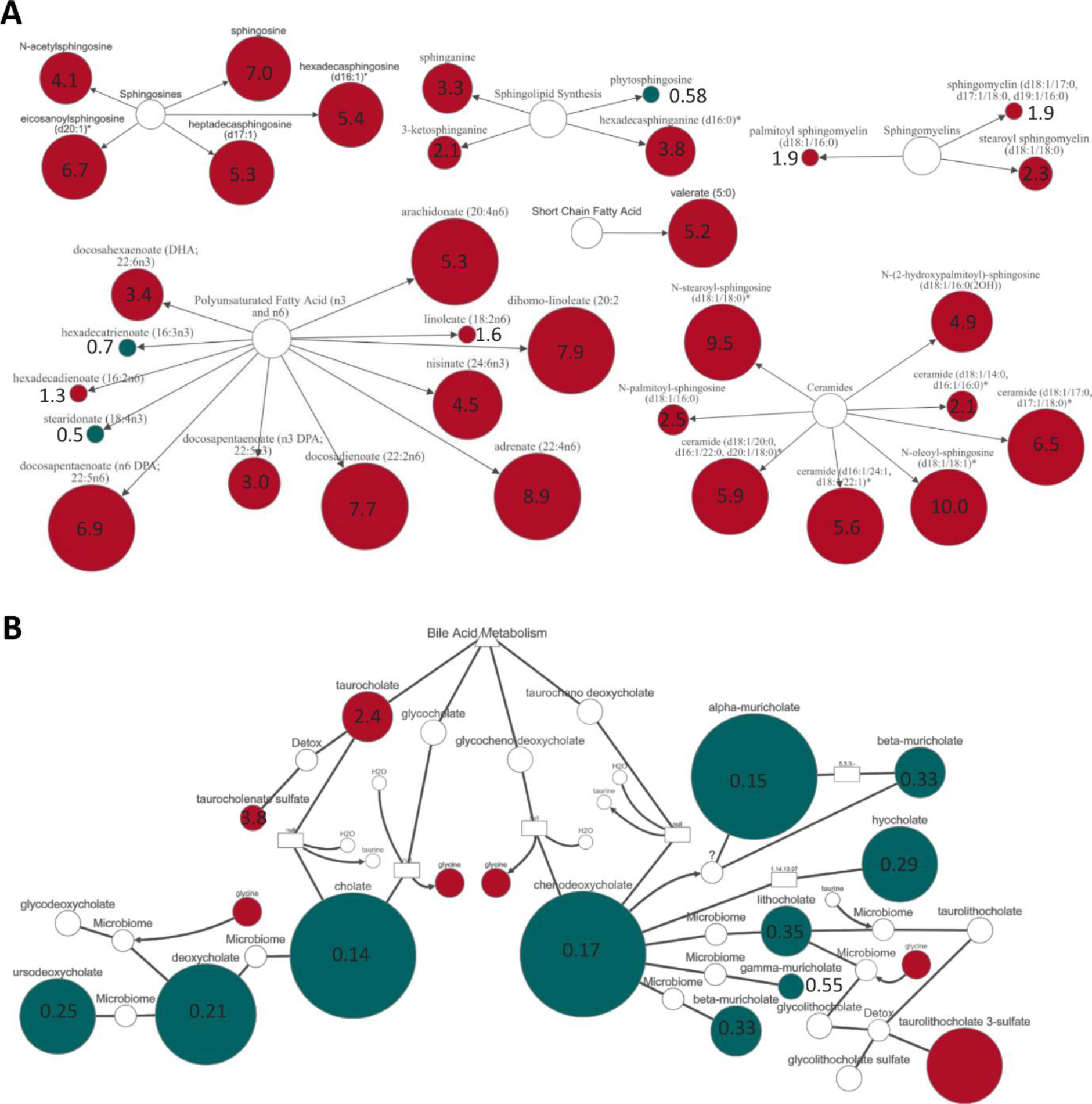
Fold change of individual lipid metabolites in the faeces following RSV infection. Metabolites which significantly altered in abundance from day 0 to day 7 belonging to the sphingosine, sphingolipid, sphingomyelin, ceramide, PUFA and SCFA sub-pathways **(A)** and to secondary bile acid metabolism **(B)**. Red indicates a significant (*P* ≤ 0.05) increase in the abundance of metabolites at day 7 post RSV infection (RSV D7/D0, metabolite ratio is ≥ 1.00). Green indicates a significant (*P* ≤ 0.05) decrease in metabolite abundance (RSV D7/D0 metabolite ratio is < 1.00). Size of node indicates magnitude of fold change but is relative for each pathway. Metabolites in the sphingosine, SCFA and secondary bile acid metabolic pathways which were significant increased (red) or decreased (green) at day 7 post RSV infection compared to before infection. Size of node indicates magnitude of change in abundance.

In faecal metabolomics it can be difficult to determine whether the biochemicals detected are produced via host or microbiota metabolism because there is often cross-over between the sources. Primary bile acids however are synthesised by the liver, while secondary bile acid metabolism is conducted by gut bacteria. Primary and secondary bile acid metabolism were found to have pathway enrichment values of 1.71 and 1.36 respectively. Several primary bile acids, including cholate and chenodeoxycholate had decreased abundance at day 7 after RSV infection (Fig.6B). This corresponded with decreased abundance of the secondary bile acids deoxycholate and lithocholate. Overall the faecal metabolome was significantly different following RSV infection, with noticeable changes in lipid metabolism.

## Discussion

Other studies in mice have observed that respiratory virus infection is associated with a change in the gut microbiota, but none have proposed that infection-induced inappetence is the driver (Wang *et al*., 2014; Deriu *et al*., 2016; Bartley *et al*., 2017; Yildiz *et al*., 2018). Bartley *et al*. (2017) observed that influenza infection drives the gut microbiota composition towards a profile similar to that of calorie restricted mice. Deriu *et al*. (2016) did not observe any effect on gut microbiota composition when IFNAR^-/-^ mice were infected with influenza virus, despite the fact that the mice still lost weight after infection and that IFNAR^-/-^ mice are reported to experience more severe weight loss following respiratory viral infection (Goritzka *et al*., 2014). Yildiz *et al*. (2018) observed that the decrease in gut bacterial load coincided with weight loss after influenza infection, but they did not find any significant correlation between percentage body weight and 16S rRNA gene qPCR data. However, as the authors note, there was a discrepancy between their abundance data when measured via 16S qPCR versus 16S rRNA gene sequencing.

One question remains as to what is driving the alteration in food intake during respiratory viral infection. Our data suggest that weight change is independent of viral load as after CD8^+^ depletion and TNF-α blockade lung viral load increased without increasing weight loss mirroring findings previously reported (Graham *et al*., 1991; Rutigliano and Graham, 2004). The strongest effect on weight loss after infection was depleting CD8^+^ cells with antibody (Tregoning *et al*., 2008). Interestingly, depleting CD8^+^ cells also prevented the reduction in food intake and reversed the changes in the gut microbiota. While CD8 is predominantly present on T cells, it is also found on subsets of dendritic cells and Natural Killer cells and we did not explore the effect of CD8^+^ depletion on these cell populations (Jonges *et al*., 2001; Shortman and Liu, 2002). A role for T cells in gut microbiota changes during influenza infection has also been proposed by Wang *et al*., (2014) who linked changes with IFN-γ production by lung CD4^+^ T cells that had tracked to the gut. While we have previously observed RSV specific CD8^+^ T cells tracking to different tissues, such as the spleen, after infection (Kinnear *et al*., 2018), we did not look in the intestines in the current or previous study. It should be noted we observed no increase in lymphoid infiltration in the gut after RSV or influenza infection (Groves *et al*., 2018). This leads us to think that the effect of CD8^+^ T cells on inappetence is via a secreted factor.

This secreted factor is most likely a cytokine or chemokine produced by CD8^+^ T cells. A number of cytokines have been previously linked to anorexia, including IL-6, TNF-α, CXCL8, IL-1β, IL-18, IL-2 and IFN-γ (Sonti, Ilyin and Plata-Salaman, 1996; Langhans and Hrupka, 1999). How these cytokines supress the normal desire to eat is not well understood, especially in the context of viral infection, as most models focused on dissecting the mechanism behind infection-induced inappetence have used LPS or peptidoglycan stimulation (Langhans, 2000). There is evidence suggesting both direct and indirect effects of these cytokines on the peripheral and central nervous system, and in particular on the hypothalamus, to suppress the desire to eat (Exton, 1997; Langhans, 2000). Our current study suggests that TNF-α is not the sole factor driving weight loss after viral infection. Another contender would be IL-6, however blocking IL-6 during RSV infection has been shown to enhance weight loss (Pyle *et al*., 2017). Little is known about the role of the other cytokines in inappetence during respiratory viral infection (Pyle *et al*., 2017) although the anti-inflammatory drug indomethacin, which has been shown to prevent IL-1β-induced anorexia, did not affect food intake during influenza virus infection (Swiergiel, Smagin and Dunn, 1997). Future work should be focused on identifying the link between the induced immune response to infection and changes in sickness behaviours and the gut microbiota.

The biggest changes in the gut metabolome after RSV infection were observed in lipid metabolism. Similar changes were seen in predicted KEGG pathways associated with lipid metabolism in infants with viral bronchiolitis, and an increase in predicted sphingosine gene abundance correlated with increased *Bacteroides* in the gut microbiota (Hasegawa *et al*., 2017). This, and the PICRUSt data in our study, suggest that increased sphingosine metabolism following RSV infection may be due to a change in gut microbiota metabolism. Sphingolipid metabolite abundance has also been observed to be altered in the lung metabolome after RSV infection in mice (Shan *et al*., 2018) and in the airways of children with RSV-bronchiolitis (Stewart *et al*., 2017). It is interesting to speculate whether altered gut metabolites promote host resistance to the infection or tolerance to the immunopathology (Wang *et al*., 2016; Trompette *et al*., 2018). Wang *et al*., (2016) used fatal bacterial and viral infections to investigate the benefits of inappetence. They observed that in viral infections, reduced food intake, in particular glucose, is harmful. We looked further downstream at the effect after infection on the gut microbiota and metabolome, which due to the co-evolved nature of host and microbiota could also form part of the protective effect of inappetence during infection by altering gut microbiota metabolism. While we do not know the effect of appetite loss on disease outcome, we saw an interesting increase in a number of pro-resolving lipids in the guts at the peak of infection. Whether these have an effect on the resolution of infection remains an intriguing question worthy of further investigations.

## Methods

### Animals

Adult female BALB/c (H-2^d^) Specific Pathogen Free (SPF) mice were purchased from Charles River Laboratories UK and maintained in autoclaved Individually Ventilated Cages (IVC) under positive pressure, with a mixture of Tapvei Eco-Pure Premium Aspen Chips (Datesend) and Sizzle-Pet (1034015, LBS, UK) for bedding. Mice were housed in groups of five animals per cage, except for fasting studies where mice were housed individually to control food intake. Mice had *ad libitum* access to irradiated RM3 pellets for food (801700, SDS, UK), except for the fasting studies where a set amount of RM3 pellets were provided each day. Food intake was measured during an experiment by weighing the contents of the food holder at the same time every day and dividing the amount of food eaten by the number of mice in the cage. All mice had *ad libitum* access to reverse osmosis autoclaved water. Faeces were collected from individual mice before intervention/infection and during the time course of the experiment. Mice were placed into individual disinfected pots and one pair of sterile tweezers per mouse was used to collect faecal pellets. Pellets were stored in sterile 1.5 ml Eppendorfs at -80 °C prior to bacterial DNA extraction. Pots were thoroughly disinfected between mice. At the end of the experiment mice were culled using intraperitoneal (IP) injection with 200 µl pentobarbital, followed by cervical dislocation or exsanguination under terminal anaesthesia.

All *in vivo* experiments were performed in the same specific pathogen free (SPF) room, which was maintained on a 12-h light/dark cycle at 20–24°C with 55 ± 10% humidity, at the Imperial College London St Mary’s Hospital Campus. Experiments were conducted in accordance with the United Kingdom’s Home Office regulations under Protocol number 1. All work was approved by the Animal Welfare and Ethical Review board at Imperial College London and studies followed the Animal Research: Reporting of *In Vivo* Experiments (ARRIVE) guidelines.

### Fasting Studies

All forms of enrichment, which could have been eaten during periods of fasting, were removed. Access to food *ad libitum* was also removed and dry food pellets (RM3, SDS) were cut up, weighed and placed into glass petri dishes inside the cage. Mice were weighted every day and 1.3 g of food was placed in the dish every 24 hours for a maximum of 3 days, or until 15% weight loss was reached, whichever occurred first. Control mice were given 4.3 g food every day. Individually housed mice with *ad libitum* access to food were maintained and sampled from alongside fasted/control mice to account for the stress of solo housing.

### Respiratory Infections

Mice were anaesthetised via inhalation of isoflurane and intranasally infected with 100 µl of 2 × 10^6^ PFU/ml RSV, 100 µl 4 ×10^5^ PFU/ml A/Eng/195/2009 influenza virus or 100 µl PBS per mouse.

### RSV L Gene qPCR

RNA was extracted from lung samples using phenol (QIAzol, 79306, Qiagen) and choloform extraction and a TissueLyzer (Qiagen, Manchester, UK). RNA was converted into cDNA using GoScript™ Reverse Transcription System (Product code: A5001, Promega, UK). qPCR for the RSV L gene was performed on a Stratagene Mx 3005p (Agilent technologies, Santa Clara, CA, USA) using the following primers:

5’ GAACTCAGTGTAGGTAGAATGTTTGCA-3’, 5’-TTCAGCTATCATTTTCTCTGCCAA-3’ and probe: 5’-FAM-TTTGAACCTGTCTGAACAT-TAMRA-3’

RNA copy number per µg lung RNA was determined using an RSV L gene standard(Culley, Pollott and Openshaw, 2002).

### TNF-α depletion

Mice were IP injected with 450 µg monoclonal anti-mouse TNF-α antibody (XT3.11, BE00058, BioXCell) or 450 µg monoclonal rat IgG1 isotype control (Clone TNP6A7, BE0290, BioXCell) 4 hours before being intranasally infected with RSV. Mice were IP injected again with the same antibodies on days 2, 4 and 6 post infection.

### Increasing airway TNF-α

Mice were intranasally dosed with 3 µg in 20 µl per mouse recombinant murine TNF-α (r-TNF-α, carrier-free, 575204, BioLegend Ltd, UK) every day for 3 days. Control mice were intranasally dosed with PBS every day for 3 days.

### CD8^+^ T cell depletion

Mice were IP injected with 500 µg monoclonal anti-mouse CD8α antibody (Clone 53-6.72, BE0004-1, BioXCell) or 500 µg monoclonal rat IgG2a isotype control (Clone 2A3, BE0089, BioXCell) 24 hours before being intranasally infected with RSV. Mice were IP injected again with the same antibodies on days 2 and 5 post infection.

### Flow cytometry

Bronchoalveolar lavage fluid (BALF) was collected by flushing the lungs three times with 0.5 ml sterile PBS through the trachea. The superior right lung lobe was mashed through a cell strainer and treated with Ammonium-Chloride-Potassium (ACK) lysis buffer (10-5483, Lonza). Cells were pelleted, washed with 1% BSA 0.2 mM EDTA in PBS and incubated with Fixable Live/Dead Aqua fluorescent reactive dye (L34966, Invitrogen), anti-mouse CD16/CD32 (Fc Block, Clone 2.4G2, 70-0161-V100, Tondo Biosciences), anti-mouse CD3e FITC (clone 145-2011, 11-0031-85, eBioscience), anti-mouse CD4 PE/Cy7 (clone GK1.5, 100422, BioLegend), anti-mouse CD8a APC/H7 (clone 53-6.7, 560182, BD Biosciences). Cells were acquired on a BD Fortessa flow cytometer and gated on live CD3^+^ lymphocytes. Data was analysed on FlowJo v10.1

### 16S rRNA gene sequencing

Bacterial DNA was extracted from 30 mg faeces/mouse/time point using the FastDNA^®^ Spin Kit for Soil (116560200, MP Biomedicals). A control extraction with no sample was performed for each kit and sequenced to monitor bacterial DNA contamination within the kit components. Each sequencing run contained a negative control (nuclease-free water used for library preparation), a positive control (mock community) and a kit control which corresponded to the kit used to extract DNA from samples. The V4 variable region of the 16S rRNA gene was amplified using universal bacterial primers(Klindworth *et al*., 2013) which were uniquely barcoded for each sample (Illumina Nextera Indexes Version 2). The amplicons were purified, quantified and equimolar pooled to produce a 16S rRNA gene library as described previously(Groves *et al*., 2018). Paired-end sequencing of the 8 pM denatured library, spiked with 8 pM of PhiX, was performed using the Illumina MiSeq platform (Kozich *et al*., 2013).

16S rRNA gene sequencing data was processed using the QIIME 1.9.0 software suite (Caporaso *et al*., 2010) as outlined fully in Groves *et al*., (2018). For microbiota composition analysis, operational taxonomic units (OTUs) were clustered at 97% sequence identify using UCLUST (Edgar, 2010) and open reference clustering. Representative OTUs were picked using the SILVA 115 rRNA database. Taxonomy was assigned using the RDP classifier (Cole *et al*., 2014) and the SILVA 115 rRNA database for reference sequences. Diversity and phylogenetic analysis was conducted in R 3.3.0 (R Core Team, 2015) with RStudio (RStudio Team, 2016) using the phyloseq (McMurdie *et al*., 2013) and vegan packages (Oksanen, 2017). Beta diversity was analysed using non-metric multidimensional-scaling (NMDS) ordination on Bray–Curtis dissimilarity matrix.

### PICRUSt

PICRUSt (Phylogenetic Investigation of Communities by Reconstruction of Unobserved States) predicts the metagenome of a bacterial community of bacteria using 16S rRNA sequencing data by constructing a table of expected gene abundances for each OTU based on KEGG orthology (Langille *et al*., 2013). The 16S rRNA gene sequencing data was analysed as described above and in Groves *et al*. (2018), with the exception that PICRUSt requires a closed-reference OTU table where representative sequences are picked and taxonomy assigned using an adapted Greengenes Database for PICRUSt (DeSantis *et al*., 2006), and so a new OTU table was generated for this. PICRUSt data was analysed in STAMP (Statistical Analysis of Metagenomes Profiles).

### Faecal Metabolomics

Profiling of the faecal metabolome before and during RSV infection was performed by Metabolon (Durham, NC, USA). Samples were processed, analysed and annotated by Metabolon as described previously (Evans *et al*., 2014; Farne *et al*., 2018) using Ultrahigh Performance Liquid Chromatography-Tandem Mass Spectroscopy (UPLC-MS/MS).

Analysis was performed on scaled data, where for each metabolite/biochemical the values were scaled to set the medium equal to one, and any missing values were inputted with the minimum amount for that biochemical. Pathway enrichment analysis was used to determine which metabolic pathways contained significantly more differentially abundant metabolites following RSV infection. To calculate the pathway enrichment value for a metabolic pathway between two time points, the number of metabolites belonging to a particular pathway which were significantly altered in abundance in a pair-wise comparison (k) (relative to the overall number of detected metabolites in that specific pathway [m]) were compared to the total number of significantly altered metabolites in that pair-wise comparison (n), relative to all detected metabolites in the study (N). Pathway enrichment = (k/m)/(n/N). Metabolic networks and pathways were visualised and fold change in metabolite abundance between time points calculated using Cytoscape software (Shannon *et al*., 2003).

### Statistics

All statistical analyses were performed in Graph Pad Prism V6/8, except for the Permutational Multivariate Analysis of Variance (PERMANOVA) for differences in beta diversity which was performed in R Version 3.5.0 and statistical analysis of faecal metabolites which was calculated in Cytoscape or by Metabolon. For weight loss, a repeated measures (RM) two-way ANOVA comparing weight at day 0 with weight at each time point was conducted for each group with Dunnett’s multiple comparison test. Differences in phyla and family abundance between time points and between groups were calculated using RM two-way ANOVA with Sidak’s multiple comparison test. TNF-α levels, cell counts and percentages were testing using an ordinary one-way ANOVA, comparing the mean of each group with the mean of every other group and using Dunnett’s multiple comparison test for correction.

### Data availability

Sequencing data will be uploaded to the European Nucleotide Archive under the accession number PRJEB32774. Metadata, mapping files, OTU tables, phylogenetic trees, and codes used for analysis will be uploaded to BioStudies at EMBL-EBI.

## Acknowledgements

HG is supported by an MRC-DTP award to Imperial College. This work was supported by the European Community’s European 7th Framework Program ADITEC (HEALTH-F4-2011-18 280873) and the Wellcome Trust. We thank Leah Cuthbertson and Colin Churchward (NHLI, Imperial College London) for providing support during the 16S rRNA gene sequencing data generation. We thank Michal Przydacz for assistance with coding.

## Author contributions

HG performed experiments, analysed data and wrote the manuscript. SH analysed data. MM designed studies and wrote the manuscript. MC designed studies, analysed data and wrote the manuscript. JT designed studies and wrote the manuscript.

## Supplementary Information

**Supplementary Figure 1.**
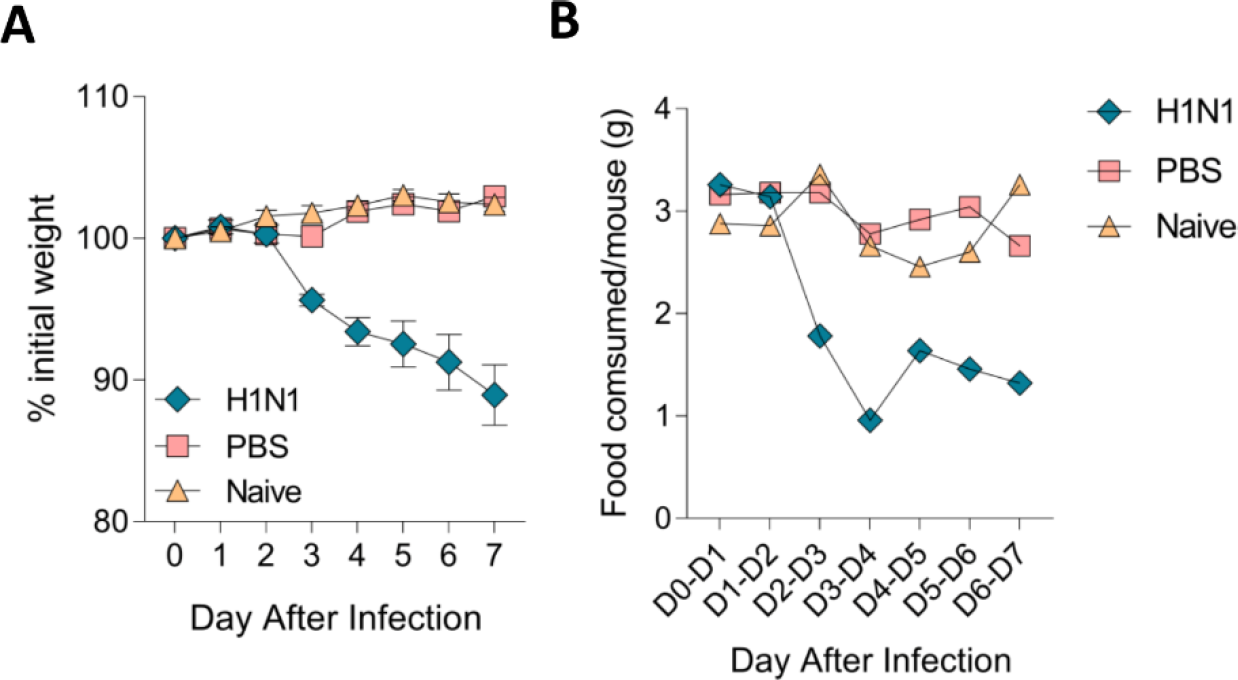
Influenza virus infection food consumption and weight loss results. Mice were infected intranasally with 4 × 10^5^ PFU/ml A/Eng/195/09 influenza virus, or dosed with sterile PBS or left naive. Mice were weighed individually **(A)** and food consumption of the entire cage **(B)** was measured every day post infection. N = 5 mice per group/cage.

**Supplementary Figure 2.**
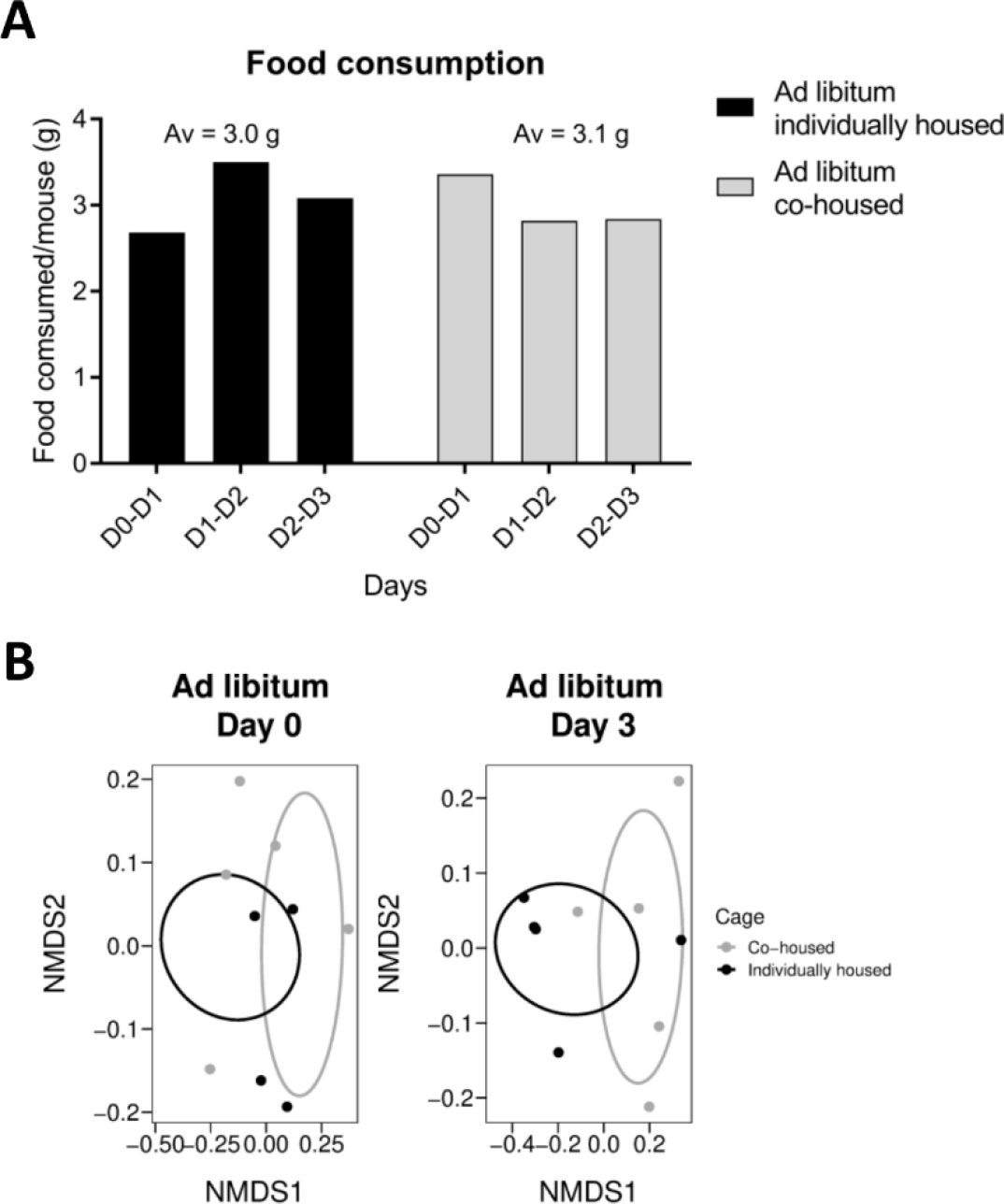
Housing individually mice does not alter food intake or gut microbiota profiles. Mice were either individually housed for 10 days or co-housed in groups of five. Food intake was measured over three days, with faecal samples taken at day 0 and day 3. N = 5 mice per group.

**Supplementary Figure 3.**
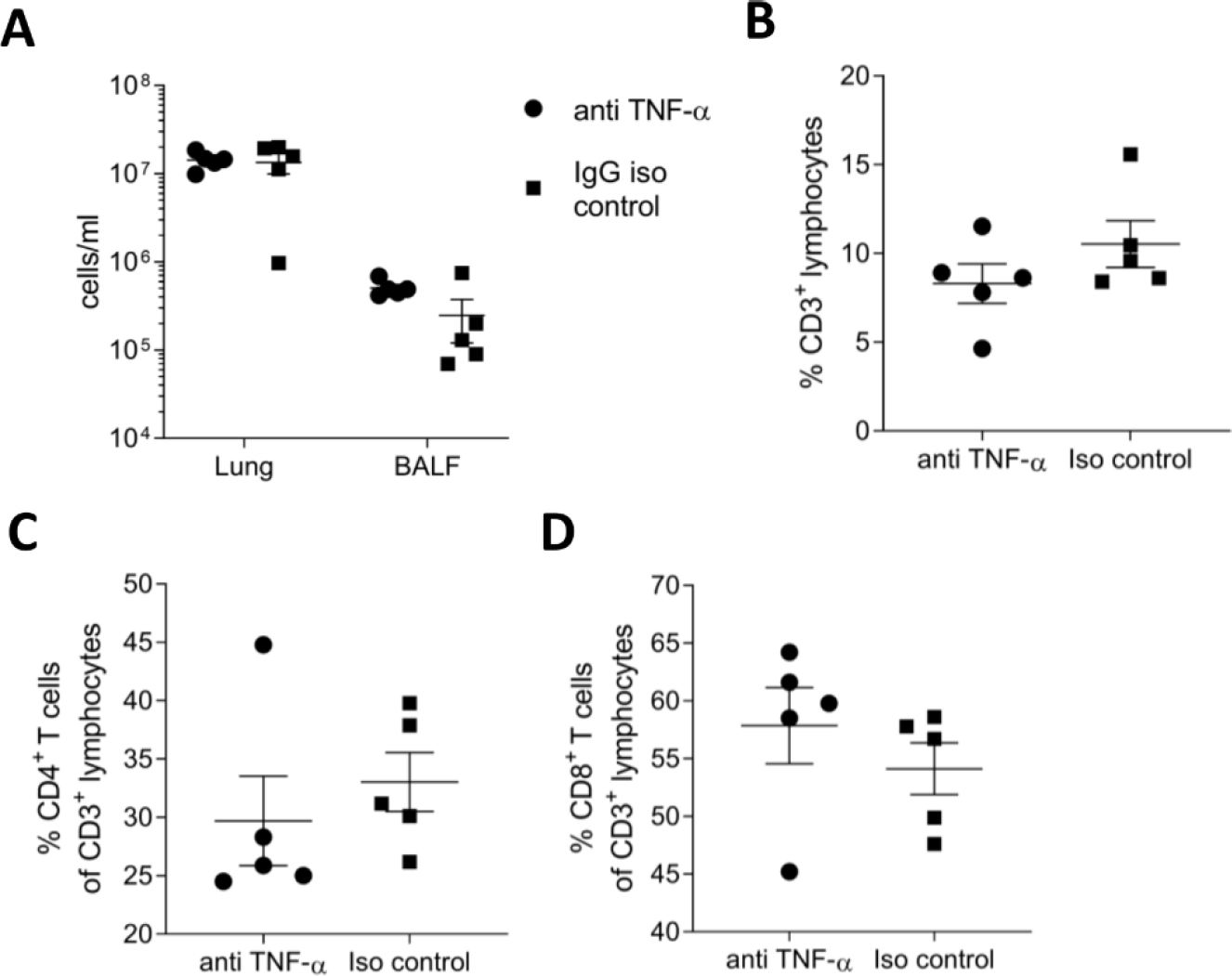
Depleting TNF-α during RSV infection increases viral load but does not alter the lung T cell response. There was no significant difference in the lung or airway (bronchoalveolar fluid [BALF]) cell counts **(A)** or in the percentage of CD3^+^ lymphocytes **(B)** CD4^+^ T cells **(C)** or CD8^+^ T cells in the lungs of TNF-α depleted mice compared to isotype control **(D)**. Unpaired t-test. * *P* ≤ 0.05. N = 5 mice per group.

**Supplementary Figure 4.**
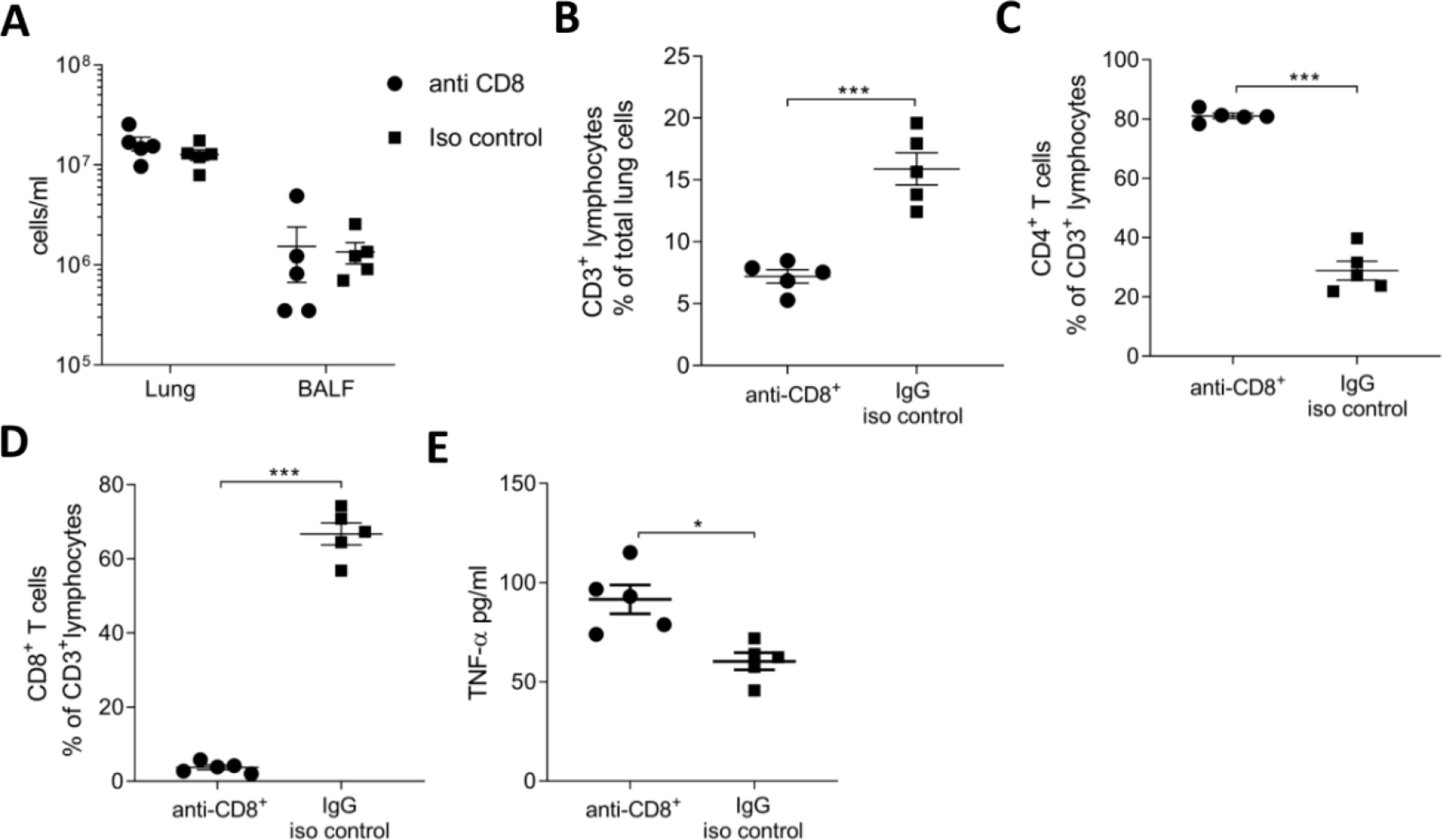
Blocking CD8^+^ T cells during RSV infection was successful and increases viral load and airway TNF-α. There was no difference in total lung or airway (BALF) cells number after blocking CD8^**+**^ T cells during RSV infection **(A)**. However, there were significantly fewer lymphocytes in the lungs of anti-CD8^+^ treated mice **(B)** and proportionally there was a much greater percentage of CD4^+^ T cells **(C)** and significantly fewer CD8^+^ T cells **(D)** in the lungs. Blocking CD8^+^ resulted in significantly higher RSV viral load **(E)** and higher airway TNF-α **(F)**. * *P* ≤ 0.05, ** *P* ≤ 0.01 *** *P* ≤ 0.001. N = 5 mice per group.

**Supplementary Figure 5:**
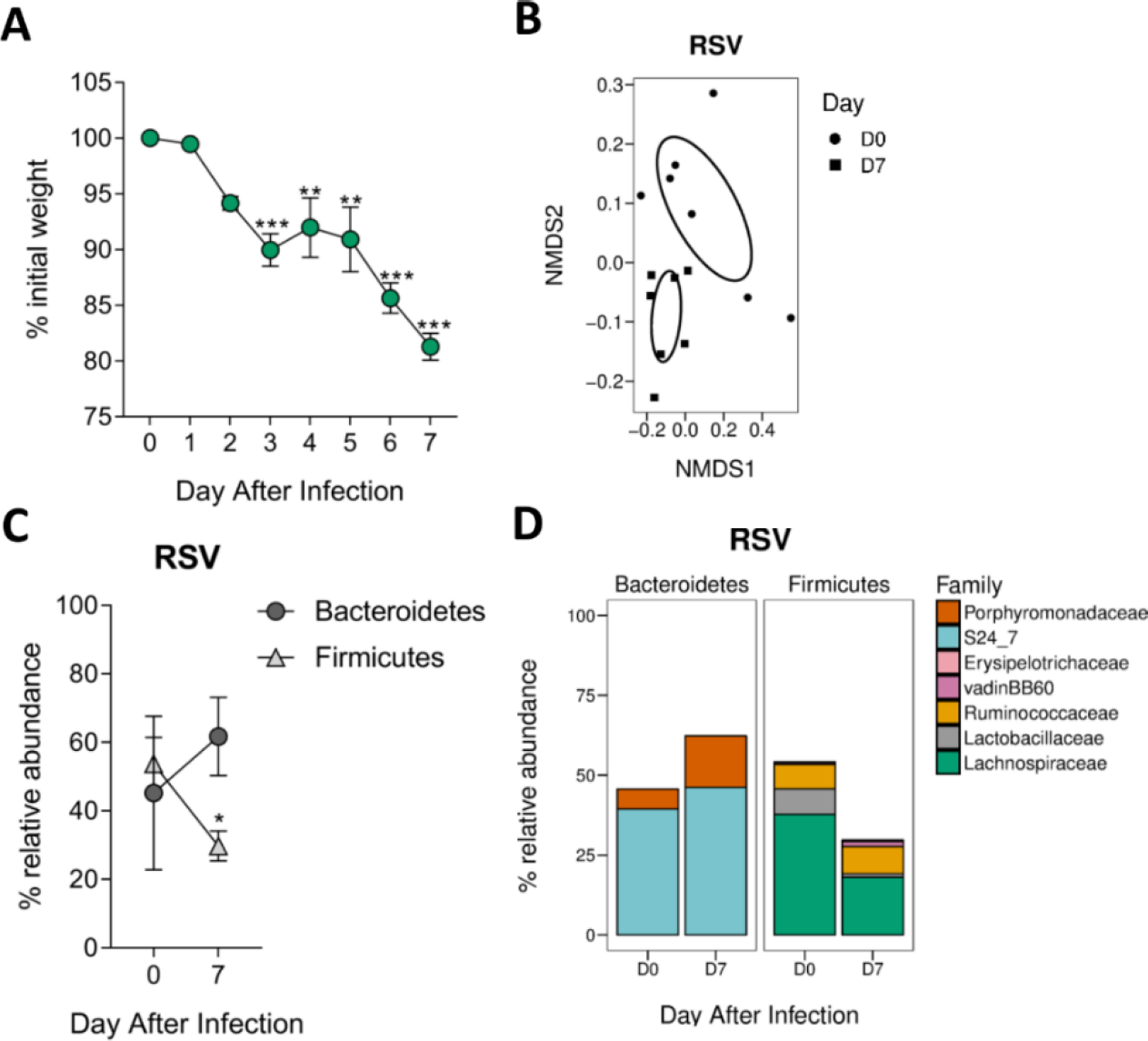
The gut microbiota is altered during RSV infection – samples matched to those used for metabolomics. Mice infected with RSV lost weight **(A)**. Diversity of the gut microbiota was significantly different after RSV infection (*P* = 0.002) **(B)**. The relative abundance of Firmicutes was significantly lower after RSV infection **(C)** with a significant increase in the relative abundance of the Porphyromonadaceae family and a decrease in the Lachnospiraceae family **(D)**. N = 7 mice. Two-way RM ANOVA with Sidak’s multiple comparison test.

**Supplementary Figure 6.**
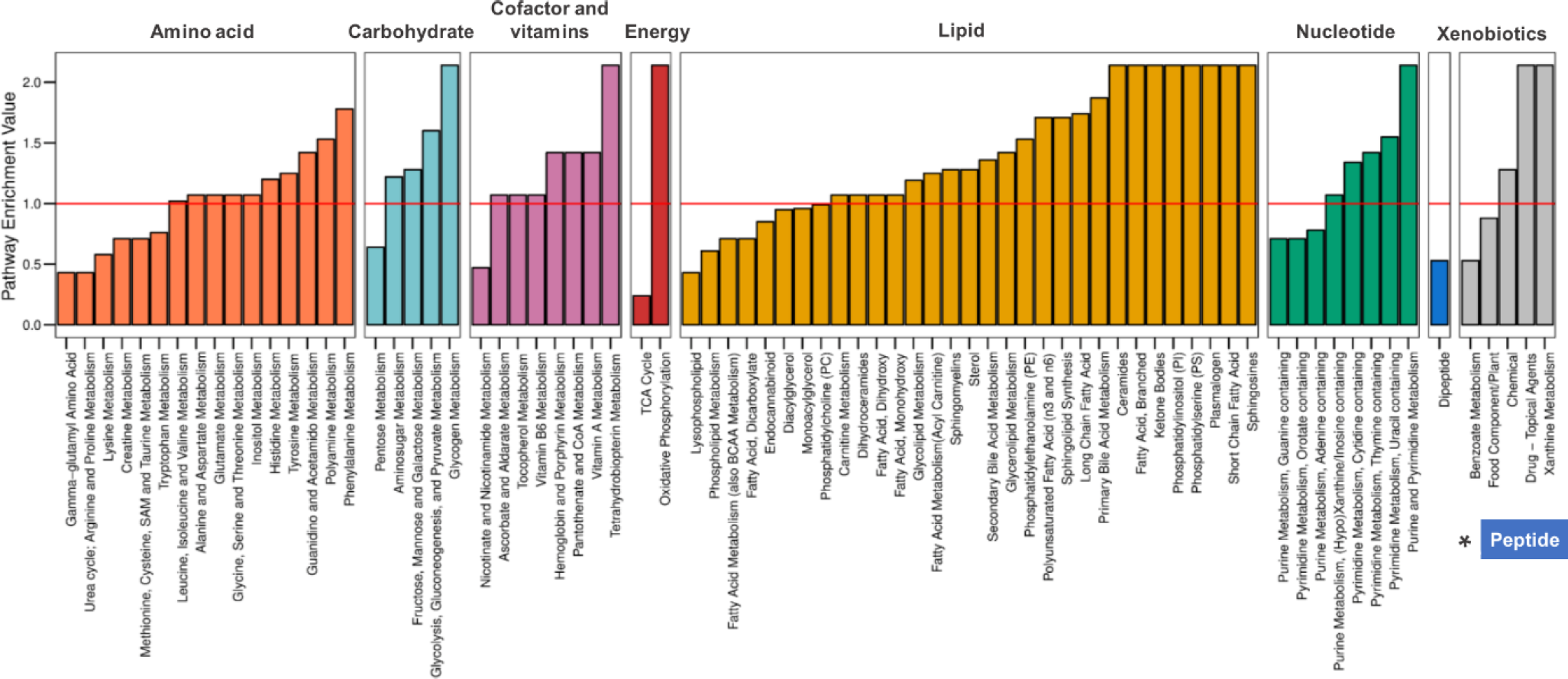
Pathway enrichment analysis of metabolite abundance comparing before infection to day 7. Pathway enrichment analysis comparing the faecal metabolome of day 7 post RSV infection samples to before infection (day 0). Red line delineates pathway enrichment values above and below 1.

